# Accuracy and reliability of [^11^C]PBR28 specific binding estimated without the use of a reference region

**DOI:** 10.1101/345645

**Authors:** Pontus Plaven-Sigray, Martin Schain, Francesca Zanderigo, Karolinska [^11^C]PBR28 study group, Ilan Rabiner, Roger Gunn, Todd Ogden, Simon Cervenka

## Abstract

[^11^C]PBR28 is a positron emission tomography radioligand used to estimate the expression of 18kDa translocator protein (TSPO). TSPO is expressed on glial cells and can function as a marker for immune activation. Since TSPO is expressed throughout the brain, no true reference region exists. For this reason, an arterial input function is required for accurate quantification of [^11^C]PBR28 binding and the most common outcome measure is the total distribution volume (V_T_). Notably, V_T_ reflects both specific binding and non-displaceable binding (V_ND_). Therefore, estimates of specific binding, such as binding potentials (e.g., BP_ND_) and specific distribution volume (V_S_) should theoretically be more sensitive to underlying differences in TSPO expression. It is unknown, however, if unbiased and accurate estimates of these measures are obtainable for [^11^C]PBR28.

The Simultaneous Estimation (SIME) method uses time-activity-curves from multiple brain regions with the aim to obtain a brain-wide estimate of V_ND_, which can subsequently be used to improve the estimation of BP_ND_ and V_S_. In this study we evaluated the accuracy of SIME-derived V_ND_, and the reliability of resulting estimates of specific binding for [^11^C]PBR28, using a combination of simulation experiments and *in vivo* studies in healthy humans.

The simulation experiments showed that V_ND_ values estimated using SIME were both precise and accurate. Data from a pharmacological competition challenge showed that SIME provided VND values that were on average 19% lower than those obtained using the Lassen plot, but similar to values obtained using the Likelihood-Estimation of Occupancy technique. Test-retest data showed that SIME-derived V_S_ values exhibited good reliability and precision, while larger variability was observed in SIME-derived BP_ND_ values.

The results support the use of SIME for quantifying specific binding of [^11^C]PB28, and suggest that VS can be used in preference to, or as a complement to the conventional outcome measure V_T_. Additional studies in patient cohorts are warranted.

## Introduction

The brain immune system has long been hypothesized to play an important role in the development and progression of neurological and psychiatric conditions (1, 2). To date, the most common method for measuring immune activation *in vivo* is to use positron emission tomography (PET) to quantify the expression of the 18kDa translocator protein (TSPO) in the brain (3). TSPO is located in glial cells, including microglia and astrocytes, and has been considered a marker for activation of these cell types (4).

[^11^C]PBR28 is a second-generation TSPO radioligand with improved signal to noise ratio (5) and reliability (6) relative to the first generation radioligand (7). It is arguably the most widely applied second-generation radioligand for examining TSPO levels in psychiatric and neurological disorders (8–10). An important goal in the field has been the evaluation of [^11^C]PBR28 as a diagnostic marker for monitoring treatment strategies that target the immune system of the brain. For this purpose, it is necessary to develop methods that provide reliable, accurate and precise estimates of outcome measures reflecting [^11^C]PBR28 specific binding to TSPO.

Since there is no region devoid of TSPO in the brain, quantifying [^11^C]PBR28 binding requires measurements of metabolite-corrected radioligand concentrations in the arterial plasma to be used as an arterial input function (AIF) in a kinetic model. When using an AIF, the most straightforward estimate of binding in the brain is the total distribution volume (V_T_), which represents the sum of the radioligand specific (V_S_) and non-displaceable (V_ND_) distribution volumes. As such, V_T_ can only be considered an indirect index of specific binding to TSPO. In contrast, V_S_ or the non-displaceable binding potential (BP_ND_=V_S_/V_ND_) are more direct estimates of specific binding (11) and should theoretically possess higher sensitivity to detect longitudinal changes or group differences. However, V_S_ and BP_ND_ calculated directly from the rate constants (estimated using a kinetic model with an AIF) are often unstable and unreliable (12,13), especially for TSPO radioligands (6, 7), and therefore of limited utility in practice.

The kinetic modelling technique Simultaneous Estimation (SIME) aims to derive a reliable, brain-wide estimate of V_ND_ in absence of a reference region (14) and consequently, more stable estimates of specific binding can be obtained. In brief, the method works by identifying the value for V_ND_ that best describes the observed PET data across all brain regions considered in the analysis. So far, SIME has been evaluated for the serotonin receptor 1A radioligands [^11^C]WAY-100635 and [^11^C]CUM101. The results showed that, for these radioligands, SIME obtained estimates that are close to “gold standard” measures of V_ND_ for these radioligands (14). With regards to [^11^C]PBR28, SIME was recently applied to quantify [^11^C]PBR28 BP_ND_ in a cohort of healthy controls and patients with Alzheimer’s disease (15). That study concluded that SIME appeared to be useful for quantification of [^11^C]PBR28, because V_ND_ and BP_ND_ were considered clearly identifiable and fell within ranges that were expected based on theory and previous publications. However, it still remains unclear whether [^11^C]PBR28 VS or BP_ND_ derived using SIME is unbiased and reliable, as the method has not yet been evaluated in cases for which the true TSPO binding levels were known.

The aim of this study was to evaluate the accuracy and reliability of SIME for estimating [^11^C]PBR28 V_ND_ and specific binding. To examine accuracy, we a) performed a simulation experiment, b) compared SIME-V_ND_ to V_ND_ estimates obtained from pharmacological competition challenge, and c) compared SIME-V_ND_, V_S_ and BP_ND_ values between high affinity binder (HAB) and mixed affinity binder (MAB) subjects in a large group of healthy controls. To examine reliability, test-retest properties of SIME-derived BP_ND_ and V_S_ values were assessed using a [^11^C]PBR28 test-retest data set.

## Methods

### SIME and measures of specific binding

SIME constrains V_ND_ (i.e., K1/k2 in a 2TCM) to be the same across a set of regions of interest (ROIs). A grid of possible V_ND_ values is then evaluated as follows: For each possible V_ND_, all ROIs are simultaneously fitted using a constrained 2TCM (in which K1/k2 is forced to be equal to the V_ND_ under evaluation). The corresponding residual sums of squares (RSS) across time frames and ROIs are then used to build an objective function for the purposes of determining V_ND_. The coordinate at which the objective function achieves a minimum is considered the optimal estimate of V_ND_ for that PET measurement. For a more detailed explanation of the SIME algorithm see (14).

In this study, the SIME-derived estimates of V_ND_ were subsequently used to calculate outcome measures of [^11^C]PBR28 specific binding according to

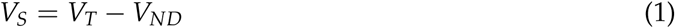

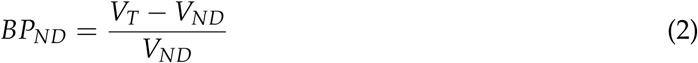

were V_T_ was independently derived from an unconstrained 2TCM in a target ROI. The primary target ROI used in this study, unless otherwise specified, is the whole grey matter defined using FreeSurfer (v5.0.0, http://surfer.nmr.mgh.harvard.edu/) segmentation.

### Subjects and data

This study includes three different datasets of healthy subjects that underwent PET examinations with [^11^C]PBR28 (Table 1). All subjects gave written informed consent prior to their participation. Their eligibility was confirmed via a health screening, evaluation of their medical history, physical and neurological examinations and routine blood tests.

**Table 1:** Demographic, genotype and radioactivity information of included datasets.

| Dataset | N | HABs | MABs | Females | Males | Age Mean | Age SD | Injected MBq Mean | Injected MBq SD |
| --- | --- | --- | --- | --- | --- | --- | --- | --- | --- |
| KI-Database | 54 | 30 | 24 | 22 | 32 | 45.2 | 17.0 | 411.0 | 58.4 |
| TestRetest | 11 | 5 | 6 | 5 | 6 | 24.5 | 3.0 | 373.4 | 61.4 |
| XBD173 Blocking | 5 | 5 | 0 | 0 | 5 | 25.2 | 7.3 | 335.6 | 7.4 |

### 1. KI [^11^C]PBR28 database

The Karolinska Institutet (KI) [^11^C]PBR28 database currently consists of 54 subjects (30 HABs and 24 MABs; 32 males and 22 females) who participated as healthy controls in a set of previously published (6, 9, 16) or ongoing [^11^C]PBR28 studies. All subjects were examined on the same PET system using identical protocols for radioligand synthesis, acquisition of transmission and emission data, and image reconstruction and analysis, as described below.

PET measurements were carried out at the PET center at KI, Stockholm, on a High-Resolution Research Tomograph (Siemens Molecular Imaging, Knoxville, TN). Individualized plaster helmets were made for each subject and used with a head-fixation device to minimize head movement during the examinations. A 6-minute long transmission scan using a single ^137^Cs source was carried out prior to each emission scan for attenuation correction. Radiosynthesis of [^11^C]PBR28 was performed as previously described (17). The radioligand was administered as a rapid bolus injected into the antecubital vein. Emission data were acquired for 75 minutes (N=19) or 90 minutes (N=35) and binned into time frames of length 8×10s, 5×20s, 4×30s, 4×60s, 4×120s, and 7×360s (N=19) or 9×360s (N=35). PET images were then reconstructed using ordered subsets expectation maximization, including modelling of the point spread function.

Arterial blood samples were acquired during the first 5 minutes of each PET examination using an automated blood sampling system (ABSS, Alogg technogies, Mariefred, Sweden). In addition, manual samples (1-3 mL) were drawn between 1 and 20 minutes post injection, in 2-minute intervals. Afterwards, manual samples were acquired in 10-minute intervals until the end of the examination. Radioactivity was immediately measured in a well counter that was cross-calibrated with the PET system. Corresponding plasma samples were obtained by centrifuging the blood samples and measuring radioactivity in the ensuing plasma using the same well counter.

Whole-blood time activity curves (TACs) were obtained by combining the ABSS and manual blood samples curves. The plasma radioactivity curve was generated by multiplying the whole-blood TAC with plasma-to-blood ratios estimated from manual plasma samples. Parent fraction of the radioligand was measured as described previously (6). To estimate the parent fraction at intermediate time points, a Hill function was fitted to the measurements and multiplied with the plasma curve to produce the final metabolite-corrected plasma curve used as AIF for each examination.

T1-weighted Magnetic Resonance Imaging (MRI) images were obtained for all subjects on a 3-T General Electric Discovery MR750 system (GE, Milwaukee, WI). ROI delineation was performed using the FreeSurfer software resulting in 12 ROIs: whole grey matter (GM), frontal cortex, temporal cortex, parietal cortex, occipital cortex, limbic lobe, thalamus, striatum, insula, anterior cingulate cortex, posterior cingulate cortex and cerebellum. All ROIs were co-registered to the corresponding PET image, allowing for extraction of regional TACs. Since a subset of subjects in the database underwent only 75 minutes of PET examination, all TACs in this study were truncated at 75 minutes to allow for consistent pooling and comparisons, unless otherwise specified.

### 2. Pharmacological competition data

Data from five healthy control subjects (all HABs, all males) who participated in a previous pharmacological competition study (18) carried out at IMANOVA Ltd London, were reanalysed to examine the correspondence between SIME-VND and VND estimates obtained from a XBD173 blocking challenge using Lassen plot. After a baseline PET, subjects received an oral dose of the selective TSPO agonist XBD173 (10 to 90mg), followed two hours later by a repeated [^11^C]PBR28 examination. Radiochemistry, imaging protocols, reconstruction, retrieval of TACs and AIF are described in the original study (18). For the present reanalysis, 9 ROI TACs were obtained from both the baseline and blocking measurement: frontal cortex, occipital cortex, temporal cortex, parietal cortex, hippocampus, amygdala, thalamus, striatum and cerebellum.

### 3. Test-Retest data

A subset of subjects (N=12) in the KI [^11^C]PBR28 database participated in a test-retest study of [^11^C]PBR28 (6). For six of them, two PET measurements were carried out on the same day, and for the other six, the PET scans were taken 2-5 days apart. One PET examination performed on a HAB subject was shortened (60 min) due to technical reasons, and this participant was therefore excluded from the test-retest analysis in this study. Image analysis and kinetic modelling for all remaining 11 test-retest subjects were carried out as described in section 1 above.

### Simulations

A simulation experiment was performed with the goal of examining whether SIME-derived V_ND_ values were accurate and precise for [^11^C]PBR28:

First, a HAB subject from the KI [^11^C]PBR28 database was randomly selected. The unconstrained 2TCM was then applied to all ROI TACs (listed above in section “1. KI [^11^C]PBR28 database”) except for the GM in order to avoid using ROIs that spatially overlap. The vascular volume fraction (vB) was fitted together with the model rate constants. From these model curves, the average K1/k2 ratio across ROIs was calculated and set to be the “true” value for V_ND_ for all ROIs in the simulation study. Noise-free curves were then calculated by defining the impulse response function for each ROI using the average value for K1/k2 with region-specific estimates of k3 and k4 (obtained from the 2TCM), and by convolving the impulse response function with the subject’s AIF. A signal corresponding to blood contribution was then added to the simulated curves, using the subject’s measured whole blood radioactivity and ROI-specific vB-values that were also estimated from the 2TCM. To maintain a realistic variance and covariance structure of noise both across ROIs and over time, a library of residuals was created from all subjects in the KI [^11^C]PBR28 database. This was done by fitting all participants’ (N=55) ROI TACs using the unconstrained 2TCM and saving all resulting residuals. From this library, residuals were then sampled and added to the simulated curves: For each time frame, the residuals from two distinct subjects were randomly selected, and a weighted sum of their residuals at this time point was added to the corresponding frame of the model curves. The weighting was performed in order to maintain the variance-covariance structure of noise between TACs (see (19) for a full explanation). This process was then repeated for all frames, resulting in one simulated noise instance. In our simulation study, 1000 noise instances were created in this way. These simulations can be conceptually interpreted as an approximation of a situation in which a single subject has been scanned 1000 times. For an in-depth explanation of the simulation procedure see supplementary material in (19). Finally, SIME was applied to each simulated noise-instance and estimates of V_ND_ were obtained and compared against the “true” V_ND_ for the underlying subject. The same procedure as described above was then applied to a MAB subject randomly selected from the KI [^11^C]PBR28 database.

In order to assess the robustness of SIME when applied to [^11^C]PBR28 we scaled the noise up by 50% in the simulated data by multiplying each residual by 1.5.

### XBD173 competition challenge

V_T_ values for each ROI (listed above in section “1. KI [^11^C]PBR28 database”) and for each subject were obtained using the unconstrained 2TCM for all baseline and blocking examinations. The revised Lassen plot (20) was applied to estimate V_ND_ for each subject separately. In addition, the Lassen plot, it has also been suggested that occupancy and V_ND_ can be estimated from a blocking data using multi-level modelling with likelihood-based techniques (21, 22). Here, we employed the Likelihood Estimation of Occupancy (LEO) (21) method to compliment to the Lassen plot. LEO has shown to produce highly accurate estimates of V_ND_ for another radioligand, but a pre-requisite of the model is that the ROI variance-covariance matrix is known. This matrix can be estimated from an independent test-retest dataset from the same radioligand. In addition to the Lassen plot, we therefore also applied LEO to the blocking data to estimate V_ND_, using the test-retest [^11^C]PBR28 examinations described above. Finally, SIME was applied to all baseline measurements, and SIME-derived V_ND_ values were compared to the outcomes from the Lassen plots and LEO. Both 70 and 90 minute TACs were used for SIME, Lassen plot and LEO, in order to examine the stability of V_ND_ over time.

### Differences between HABs and MABs

One aim of the study was to examine differences in SIME-V_ND_, and ensuing estimates of specific binding, between HAB and MAB subjects. For this goal, SIME was applied to all subjects’ ROI TACs (listed above in section “1. KI [^11^C]PBR28 database”) in the KI [^11^C]PBR28 database, using individually acquired AIFs and estimating vB for each ROI. Mean differences in SIME-derived V_ND_ values were then examined between the 32 HAB and 23 MAB subjects. VT values from GM ROI were also derived for all subjects using the unconstrained 2TCM, including estimation of vB. GM V_S_ and BP_ND_ values were calculated using equations 1 and 2. The separation between HABs and MABs using VT and SIME-derived outcomes (i.e. V_S_ and BP_ND_) was then assessed by calculating Hedges’ g effect sizes of group differences, as well as percentage differences.

In the preliminary analysis, we found an unexpected group difference in SIME-V_ND_ between HABs and MABs. Due to this, we explored potential reasons for this difference by comparing the AIF between TSPO genotype groups. This was done by calculating the area under the curve (AUC) of each subject’s AIF and by comparing the shape of the average AIF (expressed as standard uptake values, SUV) between HABs and MABs. Average AIF SUV curves for HABs and MABs were obtained by 1) aligning the peak of all curves at a selected time point and 2) subsequently taking the average of all aligned curves across subjects within each genotype group. An average input function was then calculated for HABs and MABs combined, by averaging all subjects’ centered AIF SUV curves regardless of genotype status. Subsequently, all subjects in the KI [^11^C]PBR28 database were modelled again with SIME and the unconstrained 2TCM, but this time by using the average AIF. Following this, we assessed genotype differences in resulting V_ND_, V_S_, BP_ND_ and V_T_ estimates.

### Test-Retest analysis

For the subjects in the test-retest study (6), V_T_ values from GM and SIME-derived outcomes (V_ND_, V_S_ and BP_ND_) were obtained as described above for both test and retest PET examinations. The intraclass correlation coefficient (ICC) was used as a measure of test-retest reliability; percentage average absolute variability or test-retest variability (AbsVar) was used as a measure of reproducibility; and the standard error of measurement (SEM; (23)) was used as a measure of precision. AbsVar was included for reference since it is the most common metric reported in PET test-retest studies. However, it should be noted that AbsVar scales with the additive magnitude of the outcome and is therefore not suitable for comparing absolute test-retest performance between different outcome measures.

All kinetic modelling in this study was performed in Matlab 2014 (Mathworks, Natick, MA) and all statistical analyses were performed in R (v3.3.2, “Sincere Pumpkin Patch”).

## Results

### Simulations

Figure 1 shows the results from the simulation experiment. V_ND_ estimated using SIME showed high precision and little bias for both genotypes (Panel A V_ND:True_ =1.15, mean V_ND:SIME_ =1.17 ± 0.035SD; Panel B V_ND:True_ = 0.62, mean V_ND:SIME_ = 0.63 ± 0.018SD). When amplifying the noise by 50%, SIME still provided high accuracy when estimating the ”true” V_ND_, although with somewhat lower precision (Panel C V_ND:True_ =1.15, mean V_ND:SIME_ =1.17 ± 0.082SD; Panel D V_ND:True_ = 0.62, mean V_ND:SIME_ = 0.63 ± 0.027SD).

**Figure 1:**
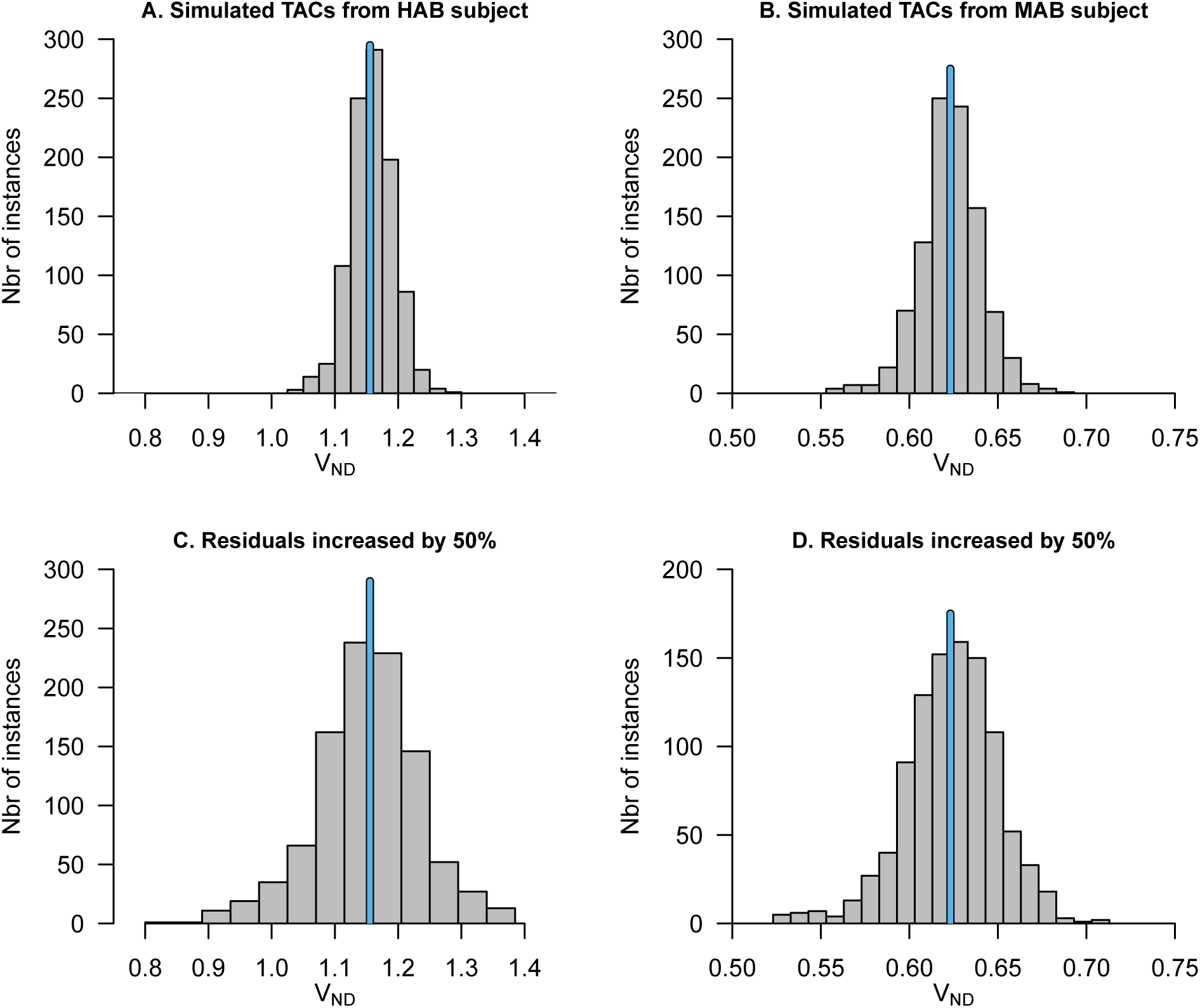
Estimation of V_ND_ from simulated time-activity curves using SIME. In each plot, 1000 noise instances have been created and added onto a set of noise-free model curves obtained from a HAB subject (A and C) or a MAB subject (B and D) randomly selected from the KI [^11^C]PBR28 database. In C and D the noise have been amplified by a factor of 1.5. The blue vertical lines indicate the “true” value of V_ND_.

### XBD173 competition

Figure 2a shows the estimated V_ND_ values from the XBD173 competition data, with 90 minute TACs, using the Lassen plot (mean V_ND_ = 1.95 ± 0.47SD), LEO (mean V_ND_ = 1.67 ± 0.51SD) and SIME (mean V_ND_ = 1.58 ± 0.37SD) methods. Figure 2b shows the estimated V_ND_ from 70 minute TACs, using the Lassen plot (mean V_ND_ = 1.72 ± 0.31SD), LEO (mean V_ND_ = 1.31 ± 0.71SD) and SIME (mean V_ND_ = 1.35 ± 0.31SD) methods. On average, SIME yielded lower V_ND_ values compared to the Lassen plot (90 min: −19%; 70 min: −22%) but displayed close correspondence to LEO (90 min: −3%; 70 min: +6%). For all three methods, estimated V_ND_ values were lower when shorter TACs were used (Lassen plot: mean −12%; LEO: mean −22%; SIME mean −15%).

**Figure 2:**
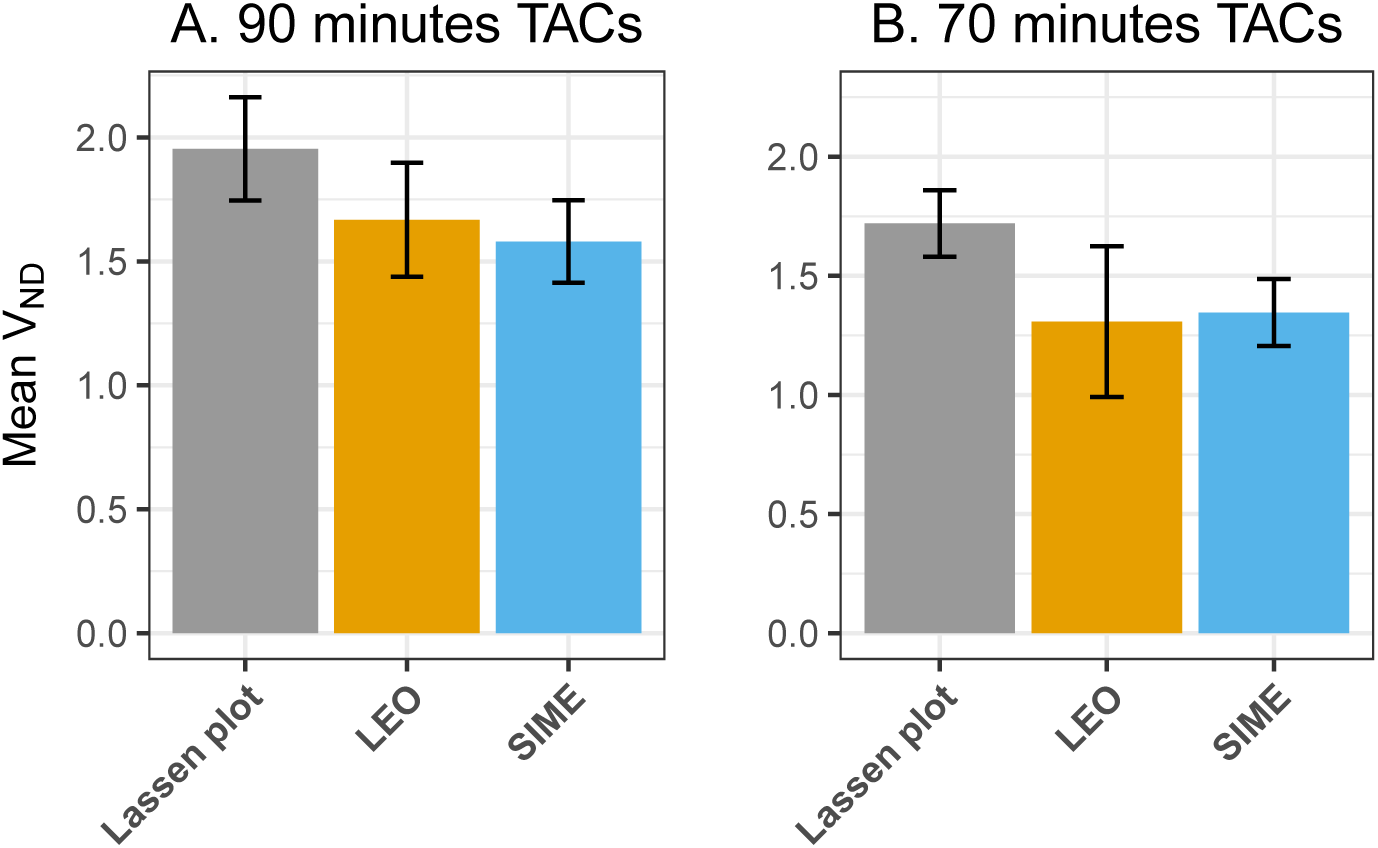
Comparison of V_ND_ estimated using different methods. V_ND_ estimates from 5 HAB subjects undergoing a XBD173 blocking challenge, using the Lassen plot, Likelihood Estimation of Occupancy (LEO), and SIME (performed only on the baseline scans). On average, V_ND_ estimated with SIME was lower than that obtained using Lassen plot, but similar to that obtained with LEO. All three methods showed lower V_ND_ when shorter time activity curves (TACs) were used.

### Differences between HABs and MABs

Figure 3a shows that there was a clear difference in V_ND_ estimated using SIME between genotype groups, with HAB subjects showing on average 35% higher V_ND_ (mean =1.31 ± 0.45SD) compared to MAB subjects (mean = 0.98 ± 0.35SD; t = 3.13, df = 52, p = 0.0028; Hedges’ g = 0.82, 95% CI [0.26, 1.38]). There was also a difference in the magnitude and shape of AIF SUV between genotype groups, with HABs showing 17% lower AUC (mean = 2365 ± 690SD) compared to MABs (mean = 2851 ± 608SD; t = −2.75, df = 52, p = 0.0083; Hedges’ g = −0.73, 95%CI [−1.28, −0.18]), and a steeper post-peak decay of radioactivity in plasma (Figure 3c and 3d). When using the same normalized AIF for all subjects, V_ND_ was similar between groups (HABs: mean = 1.01 ± 0.90SD; MABs mean = 1.03 0.7; t = −0.11, df = 52, p = 0.91; Hedges’ g = −0.029, 95%CI [-0.57, 0.51]; Figure 3b).

**Figure 3:**
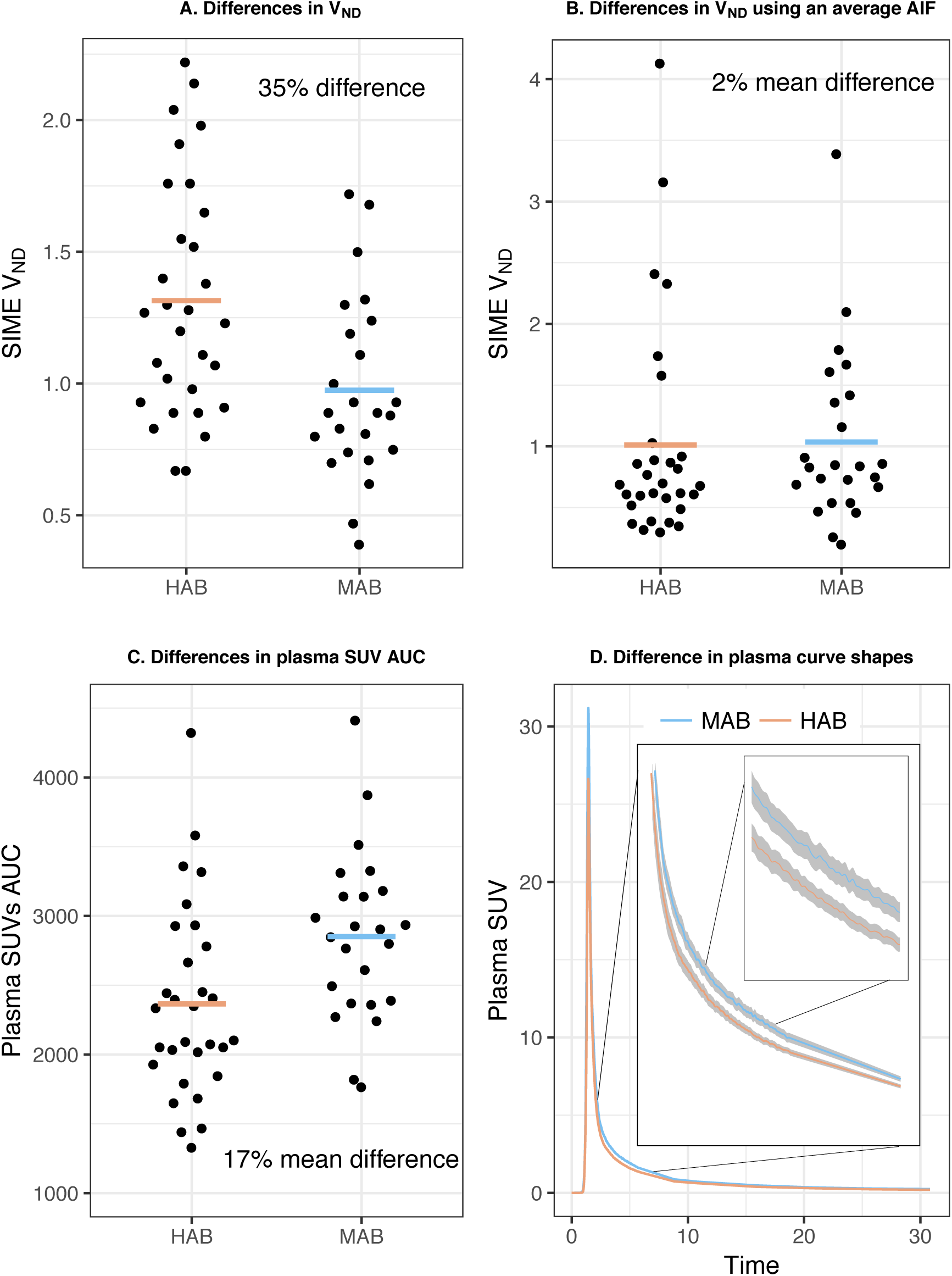
Differences in V_ND_ and input function between genotype groups. A) Difference in VND, estimated using SIME, between HAB and MAB subjects. B) The difference observed in A disappears when an average input function is used for all subjects. C) Average difference in the area under the curve (AUC) for the input function (expressed in standardized uptake values). D) Differences in peak height and shape of the average input functions between HABs and MABs. The shaded area around the lines represent 1SE. There was no overlap between the two lines in the interval seen in the subplotceptions, as determined by 95% confidence intervals.

Figure 4 shows how different outcome measures can differentiate between HAB and MAB subjects, using the GM ROI. VT from the unconstrained 2TCM (HABs mean = 4.0 ± 1.29SD; MABs mean = 2.34 ± 0.73SD; t = 6.0, df = 47, p = 2.6*10^-7^; Hedges’ g = 1.53, 95% CI [0.92, 2.14]) and VS from SIME (HABs mean = 2.69 ± 1.10SD; MABs mean =1.36 ± 0.45SD; t = 6.0, df = 40, p = 4.3*10^-7^; Hedges’ g =1.50, 95% CI [0.89, 2.11]) showed similar separation between genotype groups, while BPND from SIME (HABs mean = 2.18 ± 0.83SD; MABs mean = 1.46 ± 0.34SD; t = 4.35, df = 40, p = 8.9*10^-5^; Hedges’ g = 1.08, 95% CI [0.51, 1.66]) showed less separation. Similar results were obtained when using an average AIF for all subjects (V_T_ t = 2.41, df = 47, p = 0.02, Hedges’ g = 0.65, 95% CI [0.11, 1.21]; V_S_ t = 2.66, df = 46, p = 0.01, Hedges’ g = 0.73, 95% CI [0.17, 1.28]; BP_ND_ t =1.09, df = 51, p = 0.28, Hedges’ g = 0.28, 95% CI [-0.26, 0.82]).

**Figure 4:**
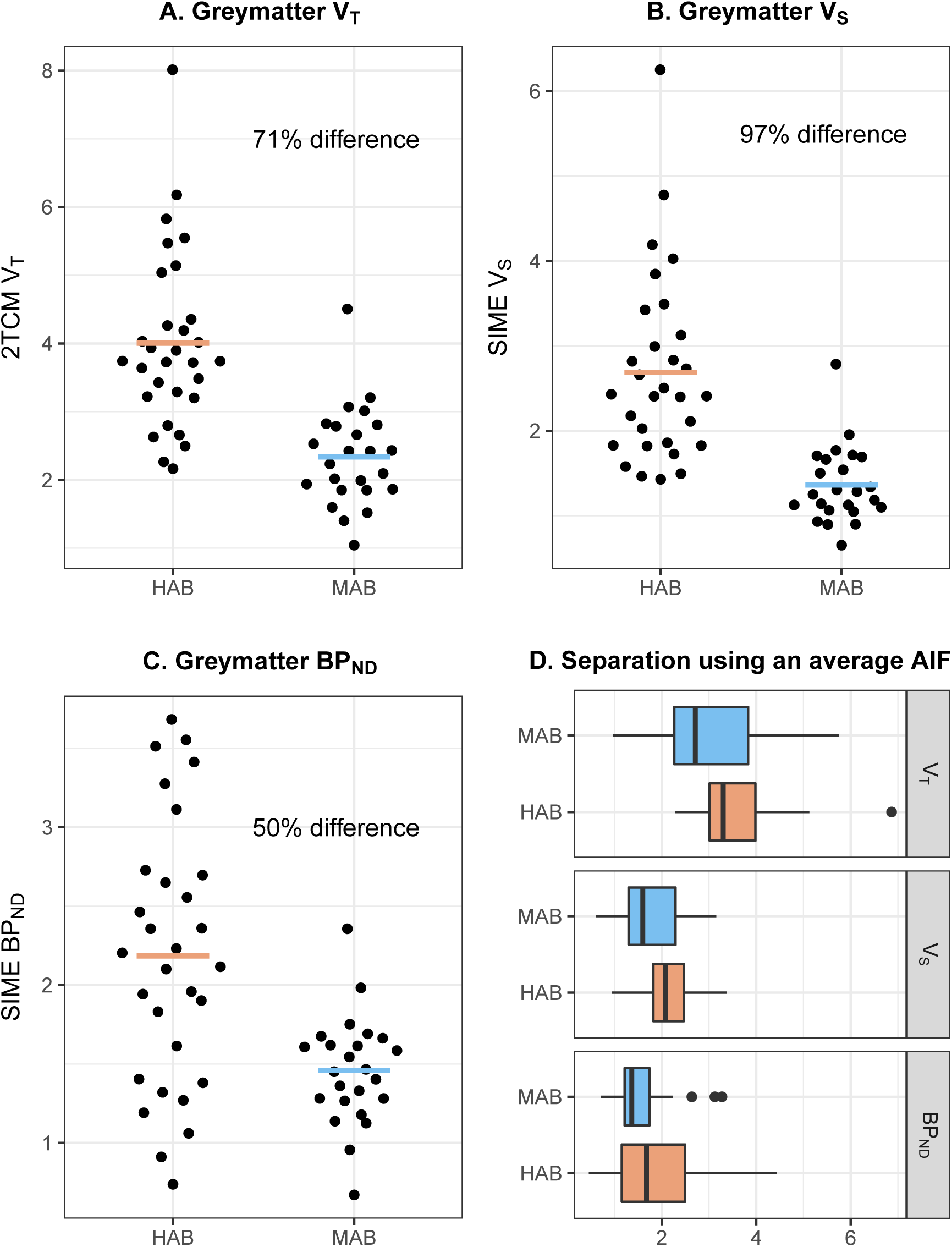
Separation of genotype groups using different outcomes. Mean percentage differences suggest that both V_T_ from 2TCM (A) and V_S_ estimated using V_ND_ from SIME (B) showed a strong separation between genotype groups, with HABs having double the VS compared to MABs. SIMEBPND showed lower mean percentage separation between HABs and MABs (C) compared to V_T_ and V_S_. When controlling for the difference in V_ND_ between genotype groups by using an average input function for all subjects, similar results were obtained (D).

### Test-Retest

Table 2 displays the test-retest metrics for all outcome measures, using the GM as ROI. VS from SIME showed excellent reliability (ICC > 0.9), while BPND showed poor reliability (ICC < 0.75) (24). VS from SIME and V_T_ from the unconstrained 2TCM showed similar reliability and precision to each other (see Table 2).

**Table 2:** Test-retest reliability, reproducability and precision estimated using the ICC, AbsVar and SEM, of different outcomes derived using 2TCM or SIME. The whole of greymatter have been used as region of interest for VT, VS and BPND. PET1 is the first examination and PET2 is the follow-up examination.

|  | PET1 Mean | PET1 SD | PET2 Mean | PET2 SD | ICC | AbsVar | SEM |
| --- | --- | --- | --- | --- | --- | --- | --- |
| 2TCM VT | 3.41 | 1.86 | 3.65 | 1.84 | 0.94 | 17.54 | 0.43 |
| SIME-VND | 1.29 | 0.47 | 1.35 | 0.47 | 0.86 | 18.07 | 0.18 |
| SIME-VS | 2.12 | 1.55 | 2.29 | 1.47 | 0.93 | 24.29 | 0.39 |
| SIME-BPND | 1.61 | 0.76 | 1.63 | 0.60 | 0.65 | 24.00 | 0.39 |

### Relationships between outcome measures

Figure 5 shows the relationship between all different outcome measures and V_ND_ values from SIME. SIME-derived V_S_ values were highly correlated with both BP_ND_ from SIME (r = 0.72, t = 7.50, df = 52, p = 7.8*10^-10^) and V_T_ from the unconstrained 2TCM (r = 0.96, t = 24.31, df = 52, p = 4.6*10^-30^). V_ND_ from SIME showed a strong correlation to V_T_ (r = 0.70, t = 7.16, df = 52, p = 2.7*10^-9^), a moderate correlation to V_S_ (r = 0.47, t = 3.88, df = 52, p = 0.00029) and no correlation to BPND (r = −0.21, t = −1.55, df = 52, p = 0.13).

**Figure 5:**
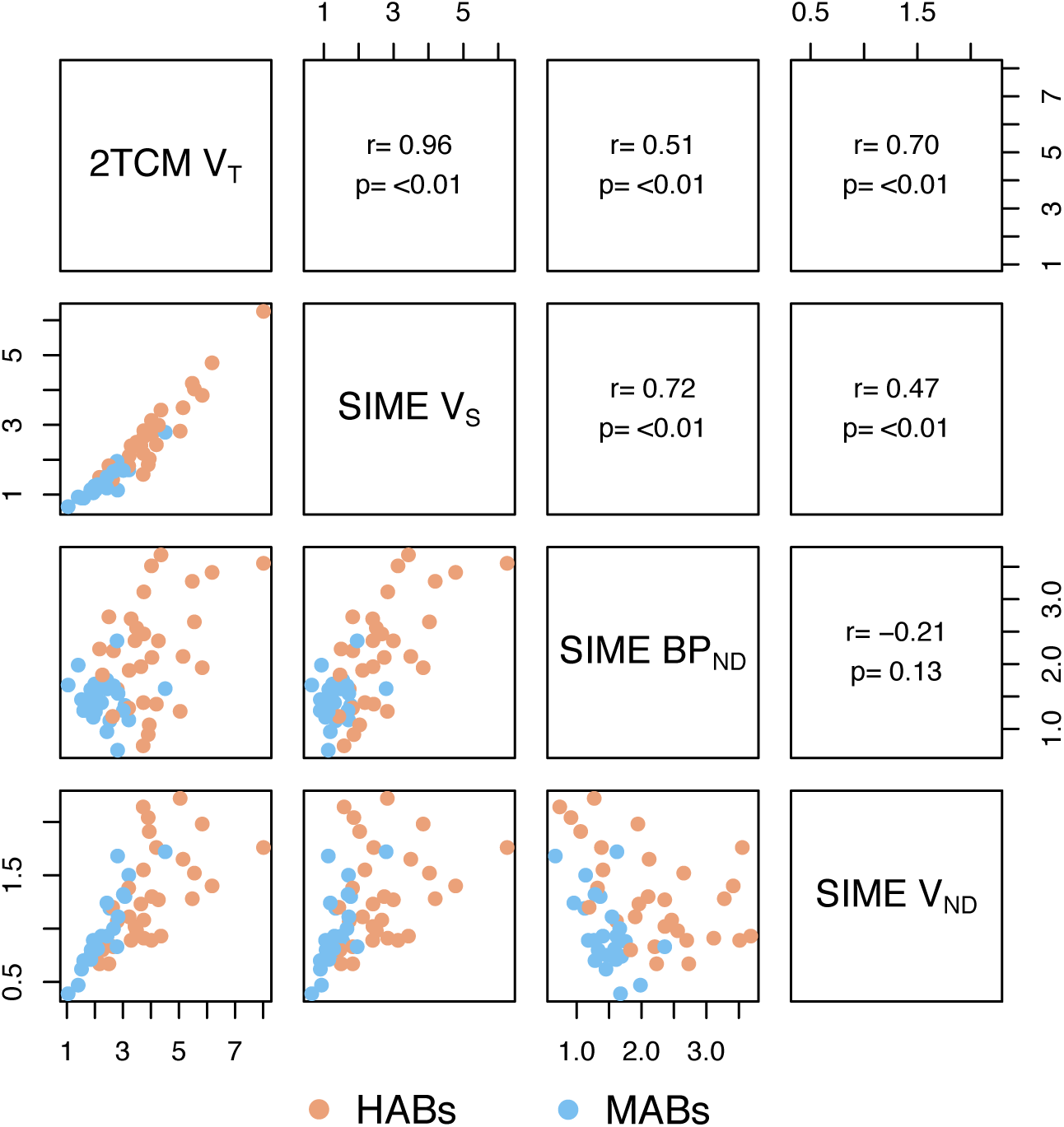
Relationships between outcome measures. Scatter plots and Pearson’s correlation coefficients (r) between V_T_ from 2TCM and V_S_, BP_ND_ and V_ND_ from SIME with the whole grey matter as region of interest.

## Discussion

Accurate, reliable and precise quantification of [^11^C]PBR28 specific binding is of high interest for clinical research, as it would theoretically lead to easier detection of effects, allowing for higher power or lower sample sizes to be used, and thereby reducing the costs in PET TSPO studies. The purpose of this study was to evaluate a new method for deriving estimates that reflect [^11^C]PBR28 specific binding (14), which has shown promising potential for [^11^C]PBR28 group comparisons (15).

We simulated [^11^C]PBR28 TACs and examined the ability of SIME to estimate a known underlying V_ND_ value. The results showed that, in simulations, SIME-derived V_ND_ values were both accurate and precise (Figure 1 A and B). This was also the case when the amount of noise in the TACs was increased above realistic levels, suggesting that SIME is robust to high levels of noise (Figure 1 C and D).

We also compared SIME against “gold standard” measures of V_ND_ by using data from a XBD173 blocking challenge (18). SIME, applied to the baseline scans, yielded V_ND_ values in the same range as the recently developed LEO technique (21), but lower than VND values obtained with the revised Lassen plot (Figure 2). Notably, for another radioligand ([^11^C]DASB) it has previously been reported that the Lassen plot tends to overestimate of V_ND,_ in particular at low occupancy levels, whereas V_ND_ estimates obtained from LEO showed to be in large unbiased, given a sufficiently powered test-retest dataset (21). It is therefore likely that the higher V_ND_ values seen with Lassen plot in the present study reflects, at least in part, inaccuracies in the Lassen plot rather than in SIME. Both the Lassen plot and LEO showed lower _VND_ when shorter TACs were analyzed, suggesting that estimates of V_ND_ are sensitive to scan duration. This trend was also reflected by the SIME method, which showed similar percentage decrease in V_ND_.

In the 2TCM, V_ND_ only reflects non-specific binding and free radioligand in tissue, which together constitute the non-displaceable binding. Since it is generally believed that the genotype only affects the radioligand’s affinity to TSPO, it follows that no difference in V_ND_ estimates between genotype groups is expected. However, in this study SIME-derived V_ND_ estimates showed a clear difference between HAB and MAB subjects (Figure 3A). We have identified three potential explanations for this observation: 1) the SIME approach is sensitive to “spill in” from the specific compartment to the non-displaceable compartment, so that SIME-derived V_ND_ values are inflated by high V_S_ values; 2) there is a systematic error in the measurement of the AIF for HABs and/or MABs that affects the estimated V_ND_; 3) a subject’s V_ND_ is dependent on the TSPO genotype (such as an TSPO affinity-dependent transport across the blood-brain barrier). To assess the first possibility, we performed additional simulations (not shown) in which the k3/k4 ratio (i.e., the BP_ND_) was both substantially increased and decreased, respectively, while the true V_ND_ was kept constant. These additional simulations showed that SIME produced similar estimates of V_ND_ regardless of the k3/k4 values, suggesting that hypothesis 1 above is an unlikely explanation to the observed difference in V_ND_ between genotype groups. As for the second possibility, we observed a clear difference between genotype groups in both AUC and shape of the plasma TAC (Figure 3C and 3D). When using a normalized input function for all subjects, the differences in SIME-V_ND_ between HABs and MABs disappeared (Figure 3B), an observation consistent with 2), but also with explanation 3) above. However, conclusions about underlying biology should not be drawn solely based on the performance of models. To date, there exists no published [^11^C]PBR28 blocking data examining V_ND_ in MAB subjects. Hence, the observed difference between genotypes cannot be fully verified, and this phenomenon warrants further investigation.

In this study, we compared SIME-derived binding values between TSPO genotype groups (Figure 4). When using individual AIFs, SIME V_S_ in HABs (mean = 2.69) was almost exactly double the value of V_S_ in MAB subjects (mean = 1.36). Assuming SIME-V_S_ is valid, this is to be expected since the low-affinity-binder allele shows negligible binding of [^11^C]PBR28 to TSPO, so that HAB subjects effectively have twice as many TSPO binding sites as MAB subjects (25).

The reliability of [^11^C]PBR28 V_S_ and BP_ND_ in GM was evaluated using a test-retest data set. SIME-derived VS showed high reliability and precision, reaching the threshold recommended for clinical use (ICC > 0.9) (24). SIME-derived BP_ND_ showed both less separation between genotype groups and lower reliability, compared to both V_T_ and SIME-derived V_S_ (Figure 4C). One potential explanation for these findings is that small amounts of measurement error in both the numerator (V_S_) and the denominator (V_ND_) of SIME-derived BP_ND_ (eq 2) leads to an amplified and larger error in the quotient, while this is not the case for subtraction carried out to calculate V_S_ (eq 1).

The results of this study supports the use of SIME-derived V_S_ as an outcome measure for future [^11^C]PBR28 examinations in preference to, or to complement, V_T_ from the unconstrained 2TCM. This is in line with the principle that V_S_ reflects more directly the level of specific binding than V_T_, and a difference of interest between subjects or groups is expected to be confined to only V_S_. For instance, if V_ND_ represents 30% of the signal, a 25% increase in V_S_ would be reflected by a 17.5% increase in V_T_, assuming both outcomes show equal variance. These hypothesized differences in sensitivity should be further tested in clinical studies using [^11^C]PBR28. To facilitate this, we publicly share all code for executing SIME in Matlab (github.com/martinschain/SIME). SIME is also implemented in the open-source R-package kinfitr for kinetic modeling of brain PET data (github.com/mathesong/kinfitr).

### Collaborators

Members of the Karolinska [^11^C]PBR28 study group are Lars Farde, Christer Halldin, Anton Forsberg, Andrea Varrone, Aurelija Jucaite, Simon Cervenka, Per Stenkrona, Karin Collste, Mats Lekander, Eva Kosek, Jon Lampa and Caroline Olgart Höglund.

## Acknowledgements

We thank David Owen, Katarina Varnäs, Anton Forsberg and Zsolt Cselénéyi for their help during the course of this investigation. We also thank the members of the Karolinska Institutet PET group, IMANOVA Ltd, Division of Molecular Imaging and Neuropathology at the New York State Psychiatric Institute and the Karolinska Schizophrenia Project for their assistance during the study.

## Author contribution

PPS conceived of the study. PPS, MS, FZ, TO and SC designed the study. TO, FZ and SC supervised the study. MS and PPS performed the image analysis and modeling. PPS, MS and SC drafted the manuscript. All authors revised the manuscript and approved the final version for publication.

## References

1. Heneka MT, Carson MJ, El Khoury J, Landreth GE, Brosseron F, Feinstein DL et al. (2015): Neuroinflammation in Alzheimer’s disease. The Lancet Neurology. 14: 388–405.

2. Doef TF van der, Doorduin J, Berckel BNM van, Cervenka S (2015): Assessing brain immune activation in psychiatric disorders: clinical and preclinical PET imaging studies of the 18-kDa translocator protein. Clinical and translational imaging. 3: 449–460.

3. Venneti S, Lopresti BJ, Wiley CA (2006): The peripheral benzodiazepine receptor (Translocator protein 18kDa) in microglia: from pathology to imaging. Progress in neurobiology. 80: 308–322.

4. Venneti S, Lopresti BJ, Wiley CA (2013): Molecular imaging of microglia/macrophages in the brain. Glia. 61: 10–23.

5. Fujita M, Kobayashi M, Ikawa M, Gunn RN, Rabiner EA, Owen DR et al. (2017): Comparison of four 11C-labeled PET ligands to quantify translocator protein 18 kDa (TSPO) in human brain: (R)-PK11195, PBR28, DPA-713, and ER176—based on recent publications that measured specificto-non-displaceable ratios. EJNMMI Research. 7: 84.

6. Collste K, Forsberg A, Varrone A, Amini N, Aeinehband S, Yakushev I et al. (2016): Test– retest reproducibility of [11C]PBR28 binding to TSPO in healthy control subjects. European Journal of Nuclear Medicine and Molecular Imaging. 43: 173–183.

7. Jučaite A, Cselényi Z, Arvidsson A, Åhlberg G, Julin P, Varnäs K et al. (2012): Kinetic analysis and test-retest variability of the radioligand [11C](R)-PK11195 binding to TSPO in the human brain - a PET study in control subjects. EJNMMI research. 2: 1.

8. Lagarde J, Sarazin M, Bottlaender M (2017): In vivo PET imaging of neuroinflammation in Alzheimer’s disease. Journal of Neural Transmission. 1–21.

9. Collste K, Plavén-Sigray P, Fatouros-Bergman H, Victorsson P, Schain M, Forsberg A et al. (2017): Lower levels of the glial cell marker TSPO in drug-naive first-episode psychosis patients as measured using PET and [11C]PBR28. Molecular Psychiatry. 22: 850–856.

10. Fregonara PZ, Richards E, Newman L, Fujita M, Niciu M, Machado-Vieira R et al. (2017): Unmedicated major depressive disorder is associated with neuroinflammation. Journal of Nuclear Medicine. 58: 139.

11. Innis RB, Cunningham VJ, Delforge J, Fujita M, Gjedde A, Gunn RN et al. (2007): Consensus nomenclature for in vivo imaging of reversibly binding radioligands. Journal of Cerebral Blood Flow and Metabolism. 27: 1533–1539.

12. Slifstein M, Laruelle M (2001): Models and methods for derivation of in vivo neuroreceptor parameters with PET and SPECT reversible radiotracers. Nuclear Medicine and Biology. 28: 595–608.

13. Varnäs K, Varrone A, Farde L (2013): Modeling of PET data in CNS drug discovery and development. Journal of pharmacokinetics and pharmacodynamics. 40: 267–279.

14. Ogden RT, Zanderigo F, Parsey RV (2015): Estimation of in vivo nonspecific binding in positron emission tomography studies without requiring a reference region. Neuroimage. 108: 234–242.

15. Schain M, Zanderigo F, Ogden RT, Kreisl WC (2018): Non-invasive estimation of [11 C] PBR28 binding potential. NeuroImage. 169: 278–285.

16. Tamm S, Cervenka S, Forsberg A, Estelius J, Grunewald J, Gyllfors P et al. (2017): Evidence of fatigue, disordered sleep and peripheral inflammation, but not increased brain TSPO expression, in seasonal allergy: A [11C] PBR28 PET study. Brain, behavior, and immunity.

17. Briard E, Zoghbi SS, Imaizumi M, Gourley JP, Shetty HU, Hong J et al. (2007): Synthesis and evaluation in monkey of two sensitive 11C-labeled aryloxyanilide ligands for imaging brain peripheral benzodiazepine receptors in vivo. Journal of medicinal chemistry. 51: 17–30.

18. Owen DR, Guo Q, Kalk NJ, Colasanti A, Kalogiannopoulou D, Dimber R et al. (2014): Determination of [11C]PBR28 binding potential in vivo: a first human TSPO blocking study. Journal of Cerebral Blood Flow & Metabolism. 34: 989–994.

19. Schain M, Zanderigo F, Mann JJ, Ogden RT (2017): Estimation of the binding potential BP ND without a reference region or blood samples for brain PET studies. NeuroImage. 146: 121–131.

20. Cunningham VJ, Rabiner EA, Slifstein M, Laruelle M, Gunn RN (2010): Measuring drug occupancy in the absence of a reference region: the Lassen plot re-visited. J Cereb Blood Flow Metab. 30. doi: 10.1038/jcbfm.2009.190.

21. Schain M, Zanderigo F, Ogden RT (2018): Likelihood estimation of drug occupancy for brain PET studies. NeuroImage.

22. Naganawa M, Gallezot J-D, Rossano S, Carson RE (2017): Quantitative PET Imaging in Drug Development: Estimation of Target Occupancy. Bulletin of mathematical biology. 1–34.

23. Weir JP (2005): Quantifying test-retest reliability using the intraclass correlation coefficient and the SEM. The Journal of Strength & Conditioning Research. 19: 231–240.

24. Portney LG, Watkins MP (2009): Foundations of clinical research: application to practice, 3rd ed.

25. Owen DR, Yeo AJ, Gunn RN, Song K, Wadsworth G, Lewis A et al. (2012): An 18-kDa translocator protein (TSPO) polymorphism explains differences in binding affinity of the PET radioligand PBR28. Journal of Cerebral Blood Flow & Metabolism. 32: 1–5.

